# An optimized somatic embryo transformation system assisted homozygous edited rubber tree generation method mediated by CRISPR/Cas9

**DOI:** 10.1101/2024.03.14.585007

**Authors:** Xianfeng Yang, Qiufei Lin, Jinu Udayabhanu, Yuwei Hua, Xuemei Dai, Shichao Xin, Xiaoyi Wang, Huasun Huang, Tiandai Huang

**Author notes:** These authors contributed equally to this work (Xianfeng Yang, Qiufei Lin, Jinu Udayabhanu).

## Abstract

Previously, we have realized the CRISPR/Cas9-RNP and plasmid mediated protoplast transient transformation genome editing in the rubber tree (*Hevea brasiliensis*), but no gene editing plants were acquired due to the bottleneck of genetic transformation. In present study, antibiotic sensitivity tests against kanamycin, hygromycin and basta were analyzed for embryo screening, the results demonstrated that 10 mg/L hygromycin is the best for transformation. Then *Agrobacterium* mediated transformation of *H. brasiliensis* embryos was carried out using a pCAMBIA1300-based CRISPR/Cas9 vector targeting Phytoene desaturase gene (*HbPDS*). High-throughput sequencing of T0 generation positive embryos which were used as regeneration materials in typical transformation procedure showed that more than 90% T0 edited embryos are chimeric with a 3.2% editing efficiency. A T0 embryo with 9.8% edited cells was sliced into small pieces for one more cycle embryogenesis to produce T1 generation embryos in order to improve the ratio of homozygous embryos. Subsequently, next-generation sequencing (NGS) demonstrated that 29 out of 33 T1 embryos were edited, nearly 50% of which were found homozygous. At last, besides four chimeric plantlets with partial albino leaves, four plantlets with complete albino phenotype were regenerated from the 29 T1 generation edited embryos, among which one is a homozygous mono-allelic mutant and the other three are homozygous bi-allelic mutants. NGS demonstrated that the threshold for the proportion of edited cells with expected albino phenotype is between 70-85%. Additionally, Tail-PCR indicate that the T-DNA was inserted into different genome positions in the four homozygous edited plantlets, combined with the different genotypes are considered, the four homozygous plantlets can be confirmed as independently derived from single transformed cells. Overall, this is the first edited rubber trees with expected phenotype reported publicly, which shows the potential in genetic improvement of *H. brasiliensis* by CRISPR/Cas9 gene editing, and subculture of T0 positive transformed somatic embryos into T1 generation is proved to be an effective and necessary procedure to produce homozygous transgenic plantlets. This study presents a significant advancement in transgenic and gene editing for rubber tree.

## 1. Introduction

Rubber tree (*Hevea brasiliensis* Muell. Arg, 2n = 36) produce most of the raw material to satisfy the industrial and national defense consumption for natural rubber (NR) demand worldwide (Priyadarshan and Goncalves, 2003). With the development of world economy, the global demand for natural rubber is increasing, at the same time, challenge aimed to improve the yield and quality of rubber tree was proposed in order to meet the demand for diversified and special high-performance rubber (Rousset, *et al*., 2021). However, as a perennial tropical tall macrophanerophytes which indigenous to the Amazon Basin in South America (Onokpise, 2004), *H. brasiliensis* has a long juvenile period up to 5-6 years before flowering, highly heterozygous genome, unstable flowering rate and poor seed set which are greatly affected by the environment conditions especially in China. At present, rubber tree varieties are mainly bred by hybrid between two backbone parental cultivars for one generation, then their F1 progeny with desired phenotype will be grafted to the stock following a suite of field evaluation spanning more than 30 years (Priyadarshan, 2017). Nevertheless, the selected varieties often carry unfavorable agronomic traits, which limits the large-scale cultivation of new varieties. Especially, nearly all the backbone parental cultivars are domesticated from Wickham collections and 1981’ IRRDB wild germplasms (Chao, *et al*., 2023), such low level of genetic diversity might hinder yield improvement in rubber tree. Due to above disadvantages and such low breeding efficiency, there has not achieved substantial progress for recent decades in *H. brasiliensis* breeding, seriously restricts the creation, spread and utilization of excellent rubber tree germplasms, thus it is an essential avenue to introduce genetic engineering tools to produce novel rubber tree cultivars more rapidly than conventional methods.

CRISPR/Cas9, as the most widely used genome editing technology, is a simple, efficient, versatile and powerful breeding tool, which has been successfully applied to genetic improvement in many plants (Li, *et al*., 2022; Li, *et al*., 2023; Liu, *et al*., 2023). Although CRISPR/Cas9 has been applied extensively for plant improvement, rubber tree is lagging far behind other crop and model species. So far, only protoplasts transient transformation editing system of *H. brasiliensis* had been established in our previous research (Dai *et al*., 2021; Fan *et al*., 2020), next we are eager to prove that CRISPR/Cas9 can function in *H. brasiliensis* by generating edited seedlings with expected phenotype.

In fact, the main obstacle for generating gene editing rubber tree is the restriction of immature transformation system for rubber tree. In the past decades, *Agrobacterium*-mediated transformation is the most wildly used method in *Hevea*, and although transgenic *Hevea* seedlings were regenerated by using embryogenic calli as receptor sources (Jayashree *et al*., 2003; Blanc *et al*., 2006; Huang *et al*., 2010; Lestari *et al*., 2018), the transformation efficiency was not satisfactory due to the low level of calli-to-embryogenesis and seedling regeneration, and the explants are not available throughout the year. Alternatively, somatic embryos (SEs) have been proved to be amenable explants for *Agrobacterium* infection in *Hevea*, and can achieve a good transformation efficiency (Huang *et al*., 2015; Udayabhanu *et al*., 2022), but a major drawback is that the regenerated cells are always not the transformed cells (Niazian *et al*., 2017; Wang *et al*., 2017), thus we inferred that the transgenic seedlings are largely chimeric although there is no powerful molecular evidence. In present study, the efficiency of embryo transformation was improved by antibiotic sensitivity analysis and adopting optimum selection strategy. Furthermore, we firstly confirm that more than 90% T0 generation positive transformed embryos are chimeric. Then by cutting one positive T0 embryo into small pieces and performing one more cycle embryogenesis subculture, 33 T1 generation embryos were produced, excitedly, nearly half of the resulting T1 embryos are homozygous. At last, eight seedlings with pleiotropic phenotypes were regenerated, of which four plantlets show complete albino phenotype, next-generation sequencing (NGS) confirmed that the mutant genotypes at the targets of *HbPDS* gene were homozygous. Tail-PCR demonstrated that the four homozygous seedlings were single transformed cell-derived. These results prove that we have overcome the limiting factor of chimeric transformed embryos, and paved the road for high-efficiency transgenic and gene editing applications in rubber tree.

## 2. Materials and methods

### 2.1. Preparation of somatic embryos

Anther-derived primary embryos were generated by our previous method (Hua *et al*., 2010), and mature cotyledonary SEs with about 1.5 cm diameter size were selected from in vitro-grown cultures, which were continuously produced by our tissue culture plants production base “Innovation Base of Natural Rubber New Planting Materials” (located in Danzhou, China) throughout the year, The embryos were then cultured on MS-based embryogenesis medium (MSE, Hua *et al*., 2010) with a slight modification (4.44 µM 6-benzyladenine, 13.9 µM Kinetin (KT), 1.44 µM Gibberellic acid, 0.27 µM 2,4-Dichlorophenoxyacetic acid (2,4-D), 70 g/L sucrose, 50 ml coconut water, 1 g charcoal and 2.2 g/L phytagel) in a designed 90 mm diameter Petri dish (Hua *et al*., 2013).

### 2.2. Antibiotic sensitivity analysis

The most popular clone of *H.* brasiliensis in China, Reyan 7-33-97 (RRI - CATAS China) was selected for the study. Newly developed mature cotyledonary somatic embryos (SE) were collected from Danzhou base. The SEs were sectioned into 2.5 mm size and cultured in a selection medium, MS based callus induction medium (MSC, Hua *et al*., 2010) supplemented with 0, 50, 75, 100, 125,150, 200, 250 and 300 mg/L kanamycin. Explants were maintained at 24 °C with a photoperiod of 24 h dark and subcultured every 20 days. The number of surviving explants were counted after 15 th, 30 th and 45 th days, and also the newly developed secondary somatic embryos were calculated to fix the resistant dose of kanamycin. Three replicates with 30 sections of SEs in each replicate, were used for every single treatment. In the similar way, triplicate experiments were performed for different concentrations of hygromycin (0.5, 2, 4, 8, 10, 12, 15, 18 and 20 mg/ L) and basta (0.25, 0.5, 0.75, 1, 1.25, 1.50 and 2 mg/ L) also. After the suitable antibiotic concentrations were fixed, pCAMBIA2301, pCAMBIA1300 and pCAMBIA3301 were used as the donor vectors which provide kanamycin, hygromycin and basta resistant respectively to evaluate the real embryos transformation effects over a longer period of selection.

### 2.3. Hygromycin selection strategies

After hygromycin was determined as the suitable antibiotic for SE transformation based on the toxicity and embryogenesis potential, four groups’ selection strategy was designed to accurately evaluate the function of hygromycin. Group 1 (G1) adopted 10 mg/L hygromycin for all the culture time, Group 2 (G2) applied 10 mg/L hygromycin for callus induction, later changed into 5 mg/L for the rest of the time, Group 3 (G3) adopted 5 mg/L hygromycin for all the culture time, Group 4 (G4) applied 5 mg/L hygromycin for callus induction, later changed into 10 mg/L for the rest of the time, each group using 5 embryos with three replicates. The gene editing vector pCAMBIA1300-2×35SCas9-*HbPDS* previously constructed by our group was used as the donor (Dai *et al*., 2021).

### 2.4. *Agrobacterium*-mediated SE transformation

The SEs were immersed in *Agrobacterium* (EHA105 strain) suspension which was cultured to OD600=0.45 in a flask supplemented with 100 µM acetosyringone for 8 minutes, then the flask containing the explants was placed in the middle of the sonicator to undergo 50 s sonication under 40 kHz strength, followed by another 10 minutes of incubation in the same liquid culture as above without shaking, and it was then kept in laminar airflow to dry the SEs using sterilized filter paper. After blot drying, it was placed on MSE for cocultivation at 22 °C with a duration of 84 h in dark conditions.

### 2.5. Induction of T0 generation positive embryos

After transformation, the infected SEs were placed on MS-based embryogenesis medium (MSE) for two days without any antibiotics. They were then cut into 3×3-mm pieces and placed on MS-based callus induction medium (MSC) containing 10 mg/L hygromycin and 500 mg/L timentin and kept in dark conditions at 24 °C. After about 20d, embryo pieces with newly sprouted T0 globular embryos were transferred to MSE containing the same selection agents. Two months later, a small piece around 2×2-mm section was cut off from the developed T0 resistant embryos for molecular detection to select the edited embryos, the positive resistant embryos were screened for T1 generation embryos induction.

### 2.6. Induction of T1, T2 generation positive embryos and plant regeneration

A positive T0 edited embryo was sliced into 2×2-mm pieces, then they were placed on the MSE for another round of embryogenesis to produce T1 resistant embryos, after molecular detection, those edited positive T1 cotyledonary embryos were transferred to plant regeneration medium (MSR,Hua *et al*., 2010) to generate edited seedlings. The culture conditions are 16 hours of light and 8 hours of dark per day at the temperature of 28 °C. Identically, one homozygous edited T1 embryo was used to produce the T2 generation embryos according the same procedure as T1 generation from T0.

### 2.7. Molecular detection of gene editing

To verify the mutated sequences at the target in transformants induced by CRISPR/Cas9, genomic DNA were isolated from T0, T1 or higher generation embryos and regenerated plantlets respectively using the Plant Genomic DNA isolation Kit (TianGen Biotech., Beijing, China). For T0 and T1 resistant embryos, they were preliminarily screened by Cas9 gene fragment PCR, following a second round PCR spanning the target of *HbPDS* gene against the positive samples detected in the first round PCR, the amplifications were subjected to NGS sequencing by the Hi-TOM platform developed by the manufacturer (Liu *et al*., 2019) to confirm the mutations. The PCR reaction was performed using PrimeSTAR Max DNA Polymerase (Takara Bio., Japan) with the following thermal cycling parameters: 98 °C for 1 min, followed by 35 cycles of 98 °C for 10 s, 56 °C for 5 s, and 72 °C for 10 s, and the final extension was performed at 72 °C for 1 min. The specific primer pairs for each amplification were shown in Table 3.

### 2.8. Tail-PCR

In order to determine the T-DNA insertion sites in the genome of regenerated homozygous editing plantlets, Tail-PCR was carried out as described by Liu and Chen (Liu and Chen, 2007). Nested primers specific to T-DNA’s Nos terminator sequence adjacent to RB and the degenerate primers are shown in Table 3. After the third round Tail-PCR, the DNA fragments were separated by 1.0 % agarose gel electrophoresis (1 × TAE buffer) and purified from the gel with the DNA Gel Extraction Kit (TianGen Biotech., Beijing, China), then they were sub-cloned into pCE2 TA/Blunt-Zero vector (Vazyme, Nanjing, China) according the manufacturer’s instructions. Those PCR positive colonies were subjected to sanger sequencing and the resulting sequences were used as the queries to blast against *Hevea* genome database (http://hevea.catas.cn/home/index).

## 3. Results

### 3.1. Antibiotic sensitivity analysis

For kanamycin test, after 30 days of culture, the survival rate of explants decreased to 47.78% when treated with kanamycin at a concentration of 100 mg/L (Table 1a), the survival of explants was further reduced as the concentration increased. At 250 and 300 mg/L, all explant tissue died during exposure to kanamycin, while at lower concentrations (150 and 200 mg/L), the death of explants began with tissue yellowing and eventually led to the cessation of explant growth (Figure S1a). Based on these findings, a treatment duration of 30 days with a concentration of 100 mg/L kanamycin was chosen for the transformation experiments. This treatment is expected to reduce the survival rate of the explants to approximately 50%. Similarly, hygromycin (Figure S1b) and basta (Figure S1c) at a concentration of 10 mg/L and 1 mg/L respectively were confirmed as the optimal selection concentration for the transformation experiment (Table 1b and 1c). Moreover, 20 mg/L hygromycin exhibited a very toxic effect at 45 days.

**Table 1a.**
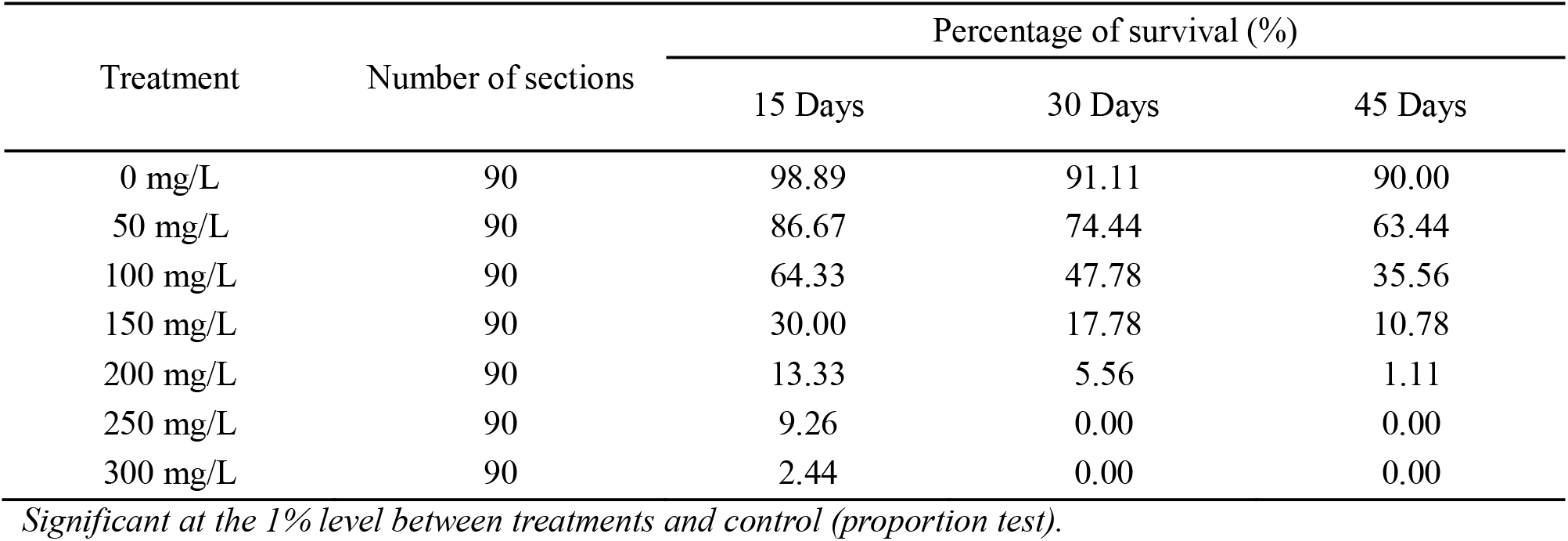
The antibiotic sensitivity tests with different concentrations of Kanamycin.

| Treatment | Number of sections | Percentage of survival (%) |  |  |
| --- | --- | --- | --- | --- |
|  |  | 15 Days | 30 Days | 45 Days |
| 0 mg/L | 90 | 98.89 | 91.11 | 90.00 |
| 50 mg/L | 90 | 86.67 | 74.44 | 63.44 |
| 100 mg/L | 90 | 64.33 | 47.78 | 35.56 |
| 150 mg/L | 90 | 30.00 | 17.78 | 10.78 |
| 200 mg/L | 90 | 13.33 | 5.56 | 1.11 |
| 250 mg/L | 90 | 9.26 | 0.00 | 0.00 |
| 300 mg/L | 90 | 2.44 | 0.00 | 0.00 |

**Table 1b.** The antibiotic sensitivity tests with different concentrations of Hygromycin. *Significant at the 1% level between treatments and control (proportion test).*

| Treatment | Number of sections | Percentage of survival (%) |  |  |
| --- | --- | --- | --- | --- |
|  |  | 15 Days | 30 Days | 45 Days |
| 0 mg/L | 90 | 100.00 | 96.67 | 95.56 |
| 0.5 mg/L | 90 | 100.00 | 92.22 | 87.78 |
| 2 mg/L | 90 | 98.89 | 90.00 | 82.22 |
| 4 mg/L | 90 | 97.78 | 87.78 | 76.67 |
| 8 mg/L | 90 | 93.33 | 66.67 | 64.44 |
| 10 mg/L | 90 | 88.89 | 57.29 | 36.67 |
| 12 mg/L | 90 | 72.22 | 33.33 | 18.56 |
| 15 mg/L | 90 | 58.89 | 23.22 | 10.00 |
| 18 mg/L | 90 | 26.67 | 12.44 | 8.89 |
| 20 mg/L | 90 | 13.33 | 9.31 | 3.33 |

**Table 1c.** The antibiotic sensitivity tests with different concentrations of BASTA.

| Treatment | Number of sections | Percentage of survival (%) |  |  |
| --- | --- | --- | --- | --- |
|  |  | 15 Days | 30 Days | 45 Days |
| 0 mg/L | 90 | 100.00 | 100.00 | 95.56 |
| 0.25 mg/L | 90 | 100.00 | 95.56 | 93.33 |
| 0.5 mg/L | 90 | 96.67 | 87.78 | 76.67 |
| 0.75 mg/L | 90 | 91.11 | 67.78 | 54.44 |
| 1 mg/L | 90 | 82.22 | 57.78 | 42.22 |
| 1.25 mg/L | 90 | 67.78 | 41.11 | 21.11 |
| 1.50 mg/L | 90 | 55.56 | 31.11 | 11.11 |
| 2 mg/L | 90 | 41.11 | 16.67 | 0.00 |

*Significant at the 1% level between treatments and control (proportion test).*

Subsequently, we carried out the transformation experiments under the determined concentrations individually using the three selection markers, it was found that both kanamycin and basta have a negative effect on growth and development of secondary resistant somatic embryos under continuous screening pressure. By contrast, hygromycin exhibits the highest potential for screening resistant embryos without sacrificing their development and growth, so 10 mg/L hygromycin was chosen as the screening marker in the initial embryo transformation.

### 3.2. Hygromycin selection strategies

The development of resistant embryos (RE) was in a different pattern in each group. The groups with constant hygromycin concentrations showed less toxicity towards the development of RE, but in G2 and G4, the growth rate was very low. It seems that the development of resistant embryos was greatly suppressed if the concentration of hygromycin was changed, G1 and G3 are better for selection because higher proportion normal REs were produced from these two groups compared to G2 and G4 (Table 2), and the positive transformed embryos were confirmed by Cas9 gene specific PCR (Fig S2). Based on the above results, maintaining a constant hygromycin concentration of 10 mg/L is the best selection strategy for embryo transformation.

**Table 2.** The results of different Hygromycin selection strategies.

| Group | Infected embryos | Resistant embryos | PCR + embryos | Percentage of PCR+ embryos to infected embryos | Percentage of PCR+ embryos to No. RE | Regenerated seedlings | Proportion of seedlings to infected embryos | Proportion of seedlings to PCR+ embryos |
| --- | --- | --- | --- | --- | --- | --- | --- | --- |
| Group1 | 15 | 36 | 21 | 140.00% | 58.33% | 3 | 20.00% | 14.29% |
| Group2 | 15 | 3 | 2 | 13.33% | 66.67% | 0 | 0.00% | 0.00% |
| Group3 | 15 | 59 | 17 | 113.33% | 28.81% | 1 | 6.67% | 5.88% |
| Group4 | 15 | 4 | 0 | 0.00% | 0.00% | 0 | 0.00% | 0.00% |

Note: PCR target gene is Cas9 gene. G1, 10 mg/L hygromycin all time. G2, 10 mg/L hygromycin for callus induction, 5 mg/L for the rest of the time. G3, 5 mg/L hygromycin all time. G4, 5 mg/L hygromycin for callus induction, 10 mg/L for the rest of the time.

**Table 3.**
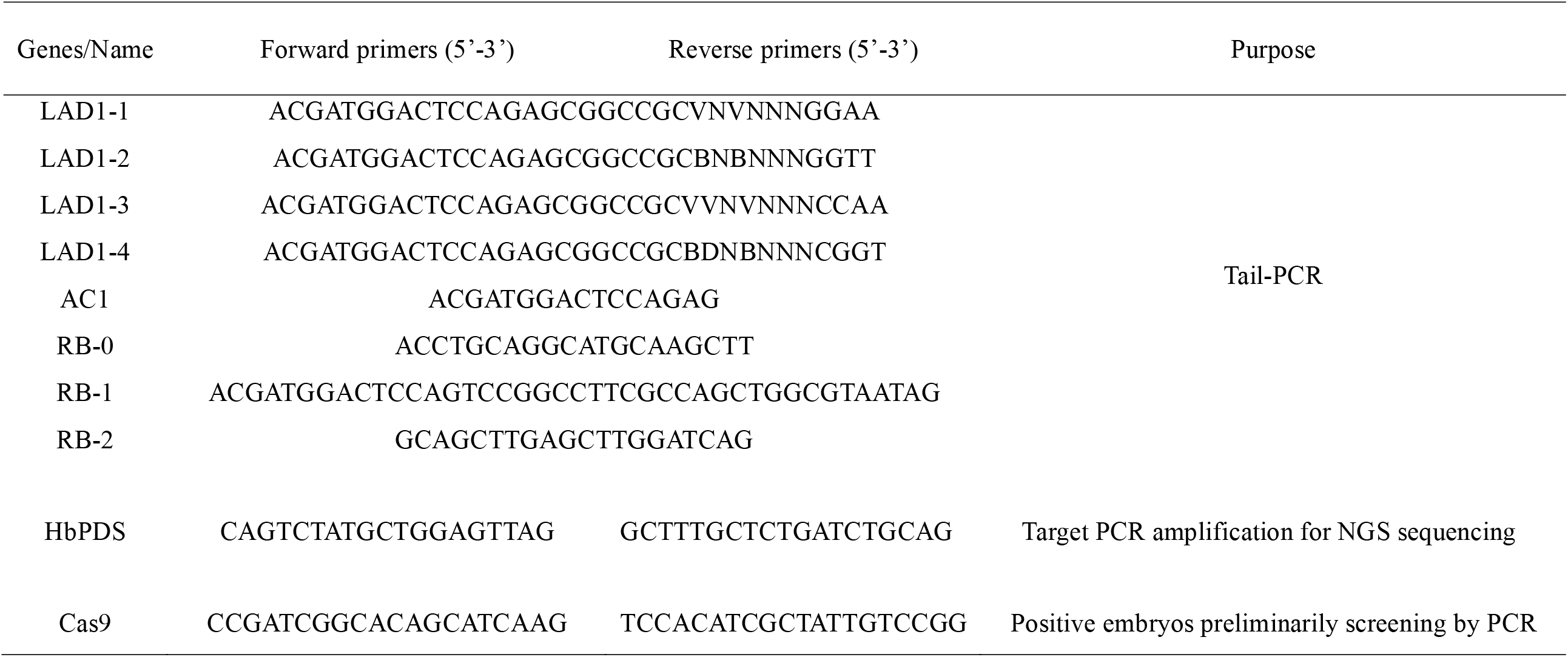
Primers used for molecular detection in present study.

### 3.3. Transformation and editing efficiency for T0 generation embryos

At the beginning, 671 T0 resistant embryos were induced by hygromycin selection from 347 *Agrobacterium* infected original embryos, among which a total of 184 embryo contained T-DNA insertions when detected by PCR amplification of Cas9 gene specific fragment (Fig S2), the result indicated that the transformation efficiency is 27.4% (184/671). Subsequently, NGS sequencing revealed that 11 of 184 Cas9 gene PCR positive embryos were edited at the *HbPDS* target, giving a 3.2% editing efficiency when counted by the initially infected 347 embryos. As shown in Table 4, the editing ratio of 11 mutated embryos span from 0.25% to 86%, 8 of them are less than or equal to 10%, 2 around 50%, only 1 above 50%. Additionally, NGS showed that there are more than three kinds of mixed mutant genotypes besides the wild sequences in each edited embryos, except E195, which comprises only single mutation type and wild type, and the proportion of both accounts for 50%. Above results suggested that, E195 was a single cell-derived heterozygous mutant, all other 10 edited T0 embryos originated from chimeric cells (90.9%), and the T0 embryos possessed relatively low editing proportion, were supposed no albino phenotype, so all T0 edited embryos were gone one more cycle secondary embryogenesis process to produce T1 generation embryos for improving the proportion of editing cells and homozygous edited embryo frequency. On the other hand, in order to test whether the T0 embryos with Cas9 gene insertion can produce target gene mutation in next generation, 62 T0 embryos with Cas9 gene insertion but without edited cells also underwent secondary somatic embryogenesis.

**Table 4.**
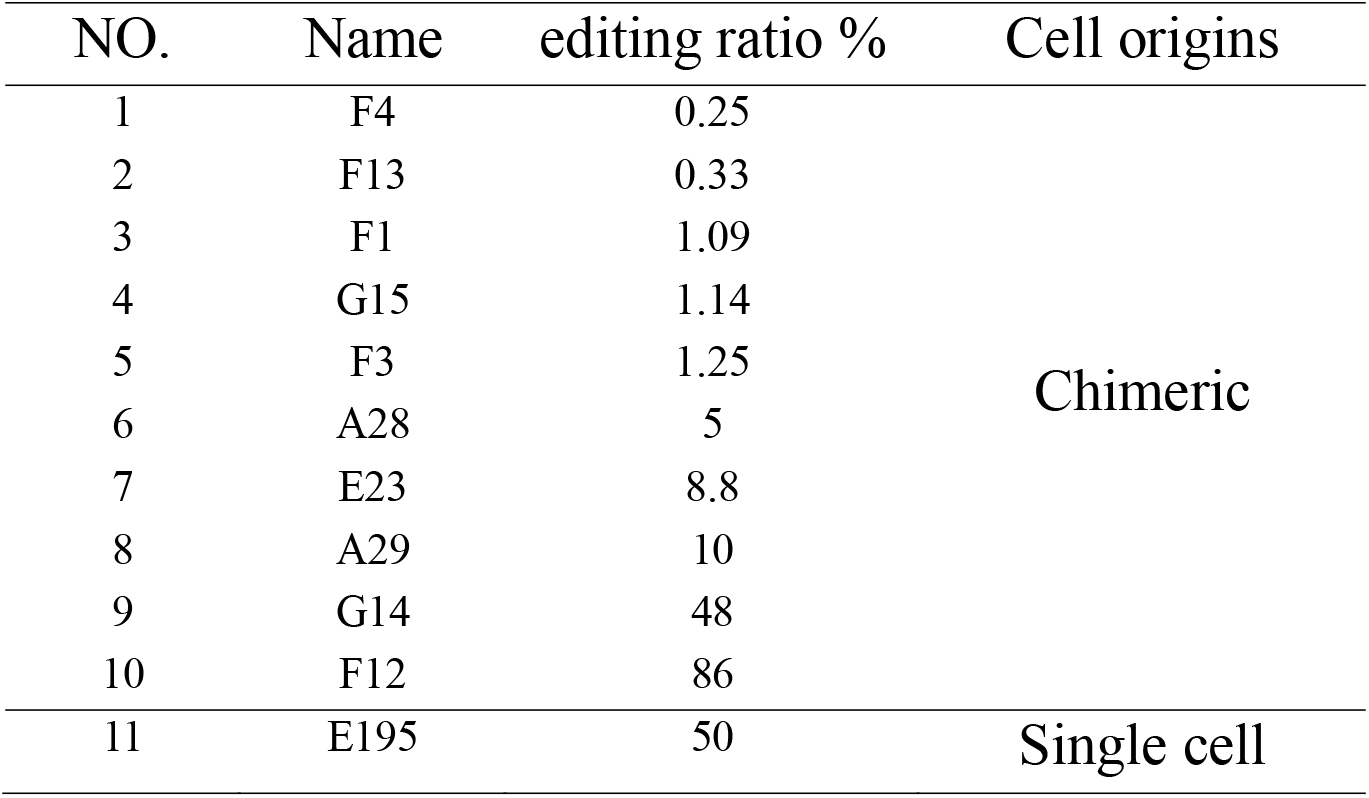
The editing ratio and cell origins of 11 T0 edited somatic embryos.

| NO. | Name | editing ratio % | Cell origins |
| --- | --- | --- | --- |
| 1 | F4 | 0.25 | Chimeric |
| 2 | F13 | 0.33 |  |
| 3 | F1 | 1.09 |  |
| 4 | G15 | 1.14 |  |
| 5 | F3 | 1.25 |  |
| 6 | A28 | 5 |  |
| 7 | E23 | 8.8 |  |
| 8 | A29 | 10 |  |
| 9 | G14 | 48 |  |
| 10 | F12 | 86 |  |
| 11 | E195 | 50 | Single cell |

### 3.4. Editing ratio and the genotypes of T1 and T2 generation embryos

3 edited T0 embryos (A29, A28 and E23) were multiplied by secondary embryogenesis successfully. A29, a T0 embryo with 9.8% edited cells composed of five types of mutation was sliced into small pieces for one more cycle embryogenesis to produce T1 generation embryos. After proliferation on the selection medium, a total of 33 resistant T1 embryos were produced, of which 29 were edited confirmed by NGS, accounting for 87.9% (Table 5). Among them, 16 embryos were single cell-derived which were composed of 13 biallelic mutants (39.4%), 2 homozygous mutants (6.1%), one heterozygous mutant (3.0%). Meanwhile, the remaining 13 embryos were chimeric, because 11 embryos contained both the edited and wild sequences, while each of the other two involved more than three kinds of mutant sequences although both of them were composed of 100% edited cells. Overall, the frequency of homozygous embryos had been improved from less than 10% in T0 to nearly 50% in T1. Similarly, A28 and E23 with only 5% and 8.8% edited cells separately were multiplied to 8 T1 resistant embryos, of which 6 T1 embryos were homozygous comprising of 1 homozygous, 2 biallelic and 3 heterozygous mutants respectively. Additionally, 20 of 62 T0 embryos with T-DNA insertion but without edited cells produced 119 T1 resistant embryos, but unfortunately, no editing case was detected, even 45 T1 embryos still contain Cas9 gene insertion. Continuously, unedited T1 embryo containing the Cas9 insertion and a homozygous T1 mutant were subcultured to T2 generation, the results showed that the progeny of the unedited T1 were still not mutated, by contrast, all the T2 embryos propagated from the homozygous T1 mutant were 100% edited with the same homozygous genotype as T1, thus we can conclude that the edited genotype can be stably inherited across generations without further segregation.

**Table 5.**
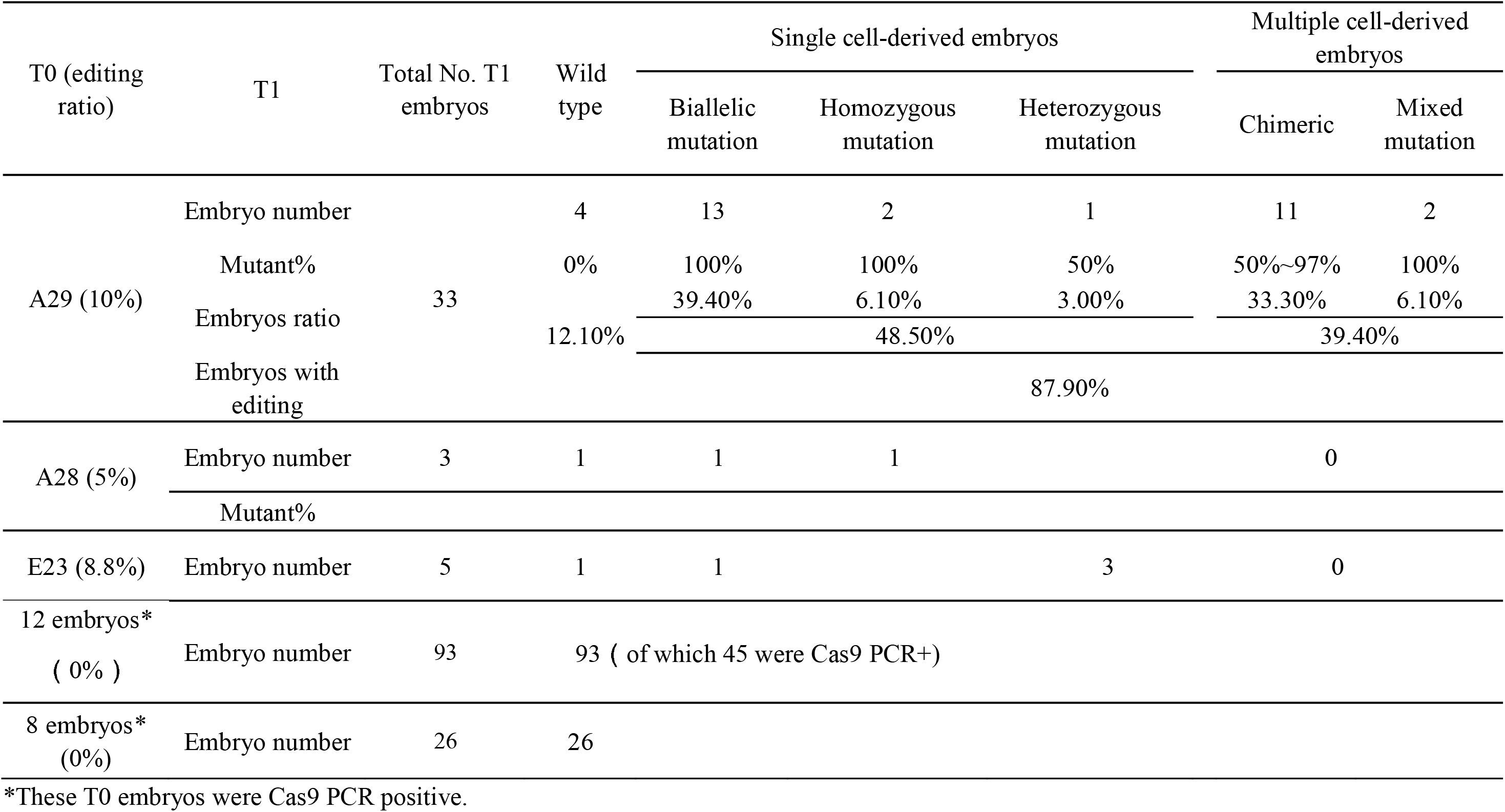
The editing efficiency and the genotype of T1 generation embryos.

### 3.5. Phenotypes and genotypes of the regenerated seedlings

Out of the 29 edited T1 embryos mentioned above, 8 gene editing plants were obtained, of which four were chimeric plants exhibiting partial albino phenotype (Designated as A to D), and four were complete albino plants (Designated as E to H, Fig. 1). We sequenced all the albino and non-albino parts (Chlorosis, pale green or mosaic) of four chimeric plants, respectively. The results showed that the proportion of mutant sequence in all non-albino parts ranges from 66% to 69%, with the only exception of the mosaic leaf in plant D, in which the proportion of wild-type sequence was only 7%, but the predominant mutation type was three-bases deletion so did not cause frameshift in the CDS of *HbPDS* gene. No wild-type sequence was detected in the albino part of plant B, and various mixed mutation types were detected. The albino part of plant C was biallelic mutation, while the mutation proportions span from 86% to 100% with various mixed mutation types in the albino leaves of plant D, demonstrating it is also a multicellular-derived chimera. Comparatively, the NGS revealed that all the four complete albino plants were homozygous with E is monoallelic mutation and F-H are biallelic mutations (Fig. 2).

**Fig. 1.**
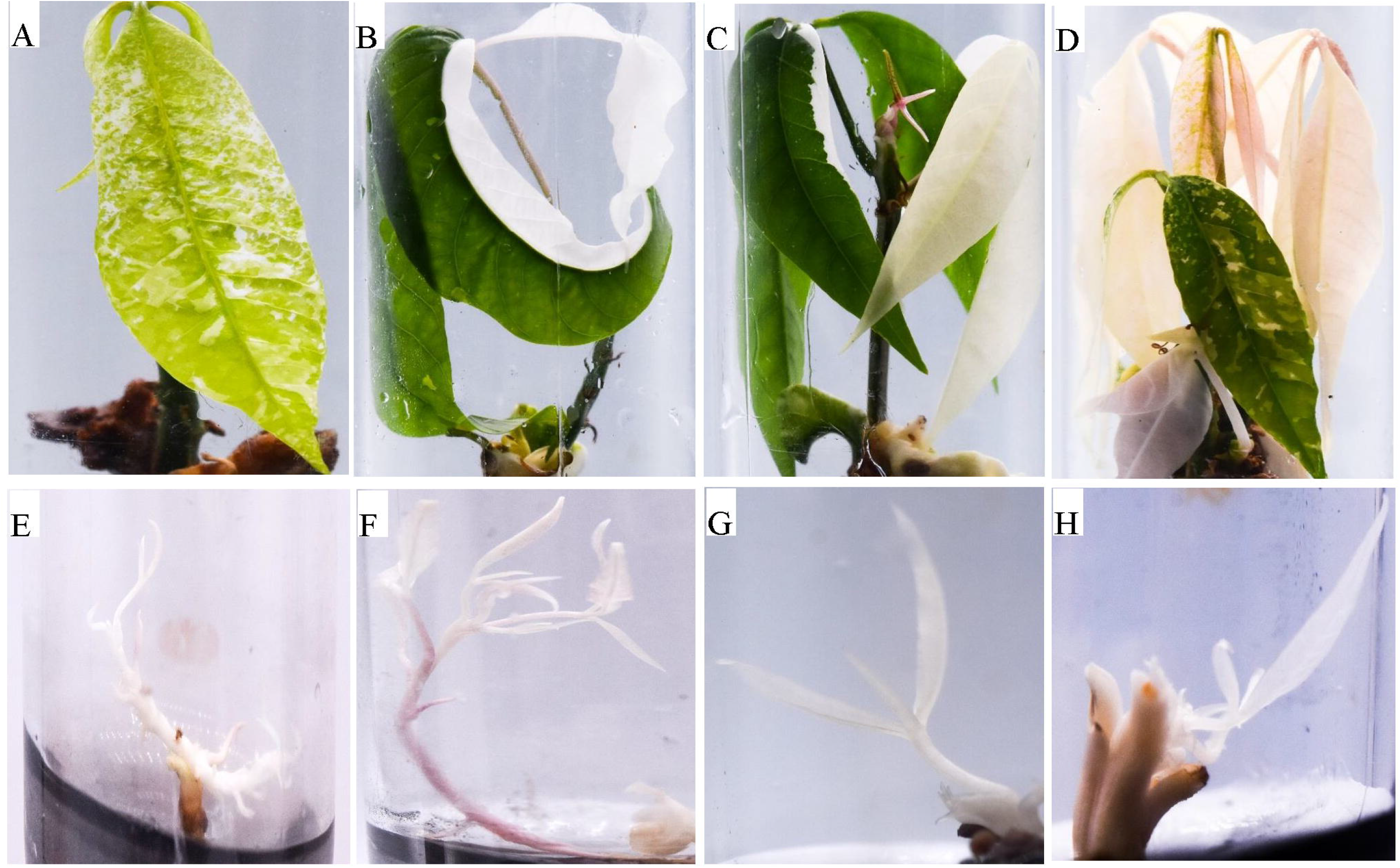
The phenotypes of *HbPDS* gene editing plantlets. A-D: Chimeric editing plantlets; E-H: Homozygous editing plantlets.

**Fig. 2.**
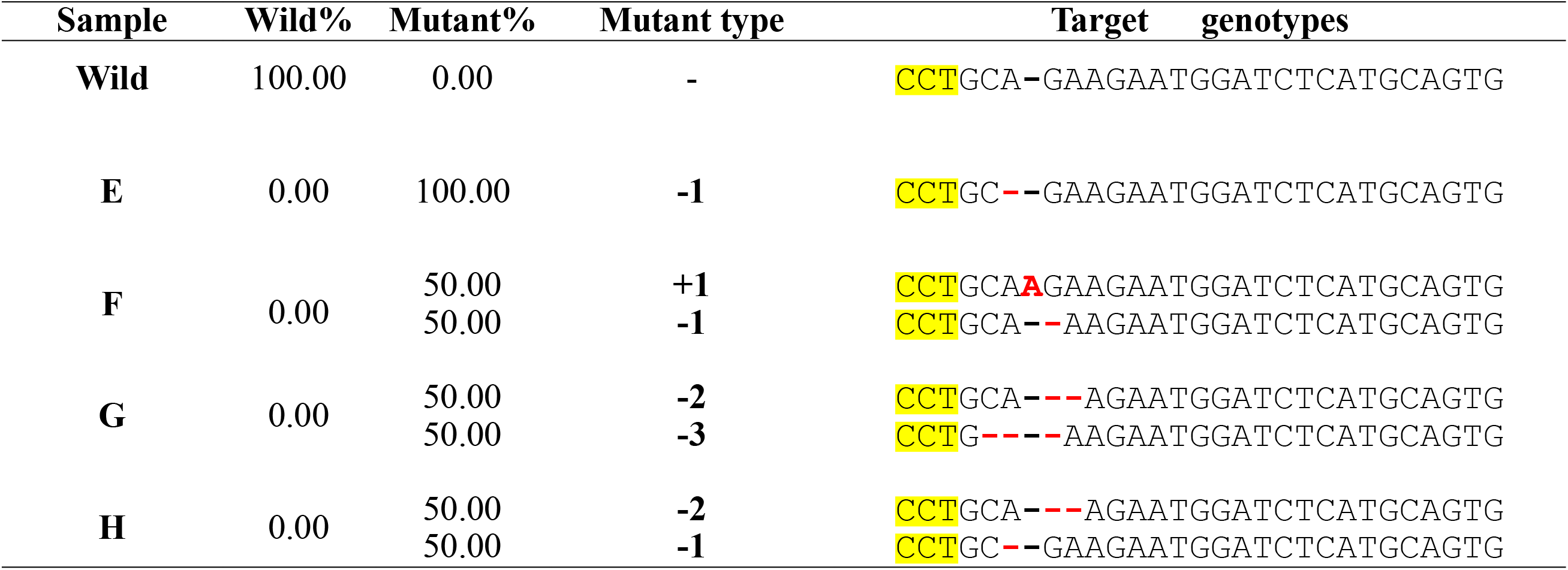
The genotypes of four homozygous edited plantlets. E is a mono-allelic homozygous mutant, F-H are bi-allelic homozygous mutants. The three yellow background letters refer to the PAM motif. The red letters indicate the mutations compared with the wild type.

### 3.6. T-DNA were inserted into different genome positions validated by Tail-PCR

The four homozygous albino plantlets generated from T1 embryos were presumed to be independently derived from single transformed cells, because the corresponding genotypes validated by NGS were different from each other. To confirm this conjecture, Tail-PCR was carried out. After three rounds of PCR, different sizes of amplicons were obtained (Fig. S3). Sanger sequencing showed that clean reads with 2376 bp, 1357 bp, 1281 bp and 1109 bp (Fig. S4), corresponding to plantlets E, F, G and H respectively were acquired after the T-DNA sequences were filtrated. Blast results demonstrated that these four sequences are aligned to different chromosome sites in the rubber tree genome (Table 6), which further confirmed that the four homozygous plantlets were developed from individual single transformed cells.

**Table 6.**
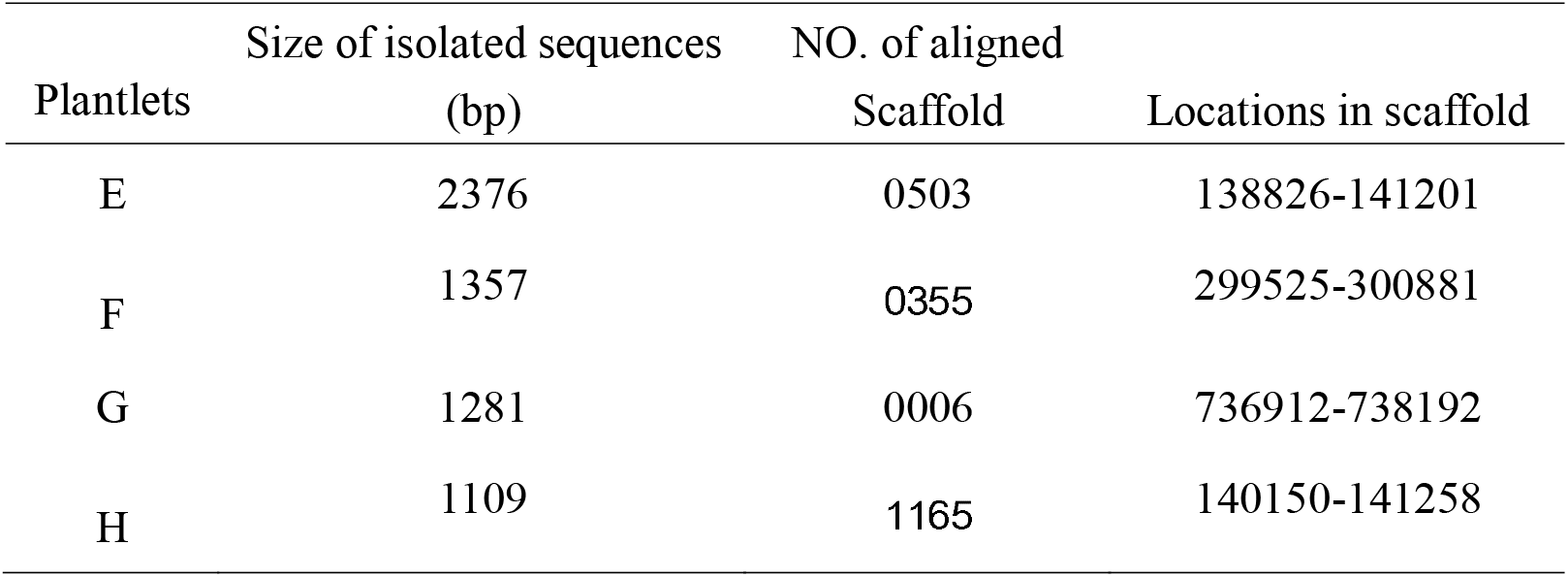
The locations of flanking sequences of T-DNA insertion in four homozygous albino plantlets.

## 4. Discussion

The genetic transformation efficiency of rubber tree is very poor compared with most plants that have established genetic transformation system, and anther-derived explants are the most widely used receptor material for genetic transformation. However, the supply of anther is affected by weather, diseases, especially florescence, resulting in the unavailable to continuously obtain receptor materials. In addition, the induction of somatic embryo from anther callus is somewhat difficult, which also impedes the genetic transformation efficiency using anther as explants. Alternatively, SEs can be used as explants to continuously provide plentiful receptor materials by highly efficient cyclic proliferation, and with relatively high transformation efficiency (Hua *et al*., 2010; Udayabhanu *et al*., 2022). Appropriate screening markers are crucial for genetic transformation efficiency, kanamycin is usually used as a suitable selection agent to screen transgenic cell lines in dicotyledonous species, including *Hevea* (Jayashree *et al*., 2003; Kala *et al*., 2014; Huang *et al*., 2015), however, the impact of different screening markers on somatic embryo transformation has not been systematically studied in *Hevea* somatic embryo transformation. In present research, we found that although there are appropriate concentrations for three antibiotics in short-term screening (less than 45 days), kanamycin and basta have strong toxic effects on embryonic growth and development over a longer duration, hygromycin was proven to be the best of the three antibiotics. Given the somatic embryo induction and development are two different process (Huang *et al*., 2015; Wang *et al*., 2017), it is presumed to increase the transformation efficiency by utilizing high concentration of hygromycin during the somatic embryo induction phase but low concentration during embryo development or shift stage. Unexpectedly, this is harmful to embryonic induction and development, and successive constant 10 mg/L hygromycin in the medium is the best favourable for embryo transformation. By combining optimized transformation methods (Udayabhanu *et al*., 2022) and suitable hygromycin screening strategy, the embryo transformation efficiency is improved from previous 4% (Huang *et al*., 2015) to 27% in present study.

In fact, it is difficult to detect whether the transformants are chimeric in the canonical overexpression transformation, because traditional PCR or qPCR tests just can determine whether the T-DNA is inserted or the target gene is expressed, but can’t differentiate transformed and untransformed cells. However, for gene editing, not only the wild and mutant cells can be detected, but also all the genotypes including wild, homozygous, heterozygous and biallelic can be thoroughly classified by NGS. Here, we clearly demonstrated that over 90% of T0 generation transformed embryos are chimeric, hence it is improper to regenerate transgenic plantlets from the T0 positive embryos. Significantly, the proportion of homozygous embryos is dramatically improved to near 50% in the T1 generation, which is consistent with the regenerated plantlets comprising four partial albino chimeric plantlets and four clear albino homozygous plantlets. It should be mentioned that all the non-albino parts in the four chimeric plantlets are also edited with 66%-69% proportion, whereas the albino parts span 86%-100% mutant rate, suggesting that the threshold for the proportion of edited cells with expected albino phenotype is between 70-85%. It can be inferred that if the T1 positive somatic embryos are subcultured to T2 or higher generation to produce regenerated seedlings, a higher proportion of homozygous plantlets will be obtained. However, considering the time cost of the subculture and the fixed genotype of higher generation progeny, it is a better choice to subculture positive T0 somatic embryos as more as possible, which can ensure the acquisition of multiple homozygous or biallelic edited plantlets with diversified genotypes. The experimental procedure in present study is simply shown in Figure S5.

Although a high proportion (27.4%) of T-DNA insertions were detected in the transformed somatic embryos, sequencing results showed that most of the positive somatic embryos did not undergo editing at the target sites. This indicates that the editing efficiency of the gene editing vector currently used is not efficient, which need to be optimized. The promoter of Cas9 gene have a significant impact not only on editing efficiency, but also have an impact on the editing type, many other promoters produce higher editing efficiency and a higher proportion of homozygous or biallelic mutations than the CaMV35S promoter (Feng *et al*., 2018; Wang, *et al*., 2022). The Cas9 gene sequence in present study is optimized based on the rice codon, driven by 2 × 35S promoter, next we will replace the 2×35S promoter of Cas9 with the endogenous constitutive strong promoter pHbUbiquitin of *H. brasiliensis* (Xin, *et al*., 2022), and optimize Cas9 gene according to the codon bias of *H. brasiliensis*.

The promoter length of U6 gene has a significant impact on the expression of driven gene. In plant gene editing system, the AtU6 promoter subsequence as short as 79 bp is used (Belhaj, *et al*., 2013). Based on the location of the two conserved elements USE and TATA box, which are located at −60 bp and −30 bp upstream of the U6 gene (Simmen, *et al*., 1990), theoretically, the promoter can exert transcription as long as these two elements are included. Previously, we also found that when the editing vector was constructed with the promoters of five *Hevea brasiliensis* HbU6 genes, the shorter promoter could lead to higher mutation efficiency (Dai *et al*., 2021). In cotton, a truncated GbU6 promoter with 105 bp size showed highest transcriptional activity than a series of longer promoters (Lei *et al*., 2016). Based on these results, we will test the editing efficiency using shorter U6 promoter. Although current gene editing vector needs to be optimized, the regeneration of four homozygous editing plantlets indicates that present research paved the way for the application of transgenic and genome editing in rubber tree.

In summary, our work contributed to improving the transgenic efficiency in rubber tree, demonstrates the feasibility to produce homozygous or biallelic edited *Hevea* plantlets, representing an important breakthrough in the application of the CRISPR technology in *Hevea* genetic improvement. Accordingly, CRISPR /Cas9 holds the potential to greatly accelerate the breeding duration of rubber tree, as well as to rapidly identify rubber tree gene functions in vivo.

