## Supplemental Figure 1 for "An optimized somatic embryo transformation system assisted homozygous edited rubber tree generation method mediated by CRISPR/Cas9"

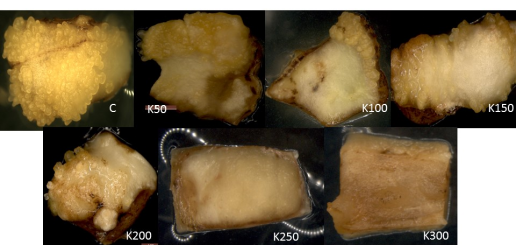

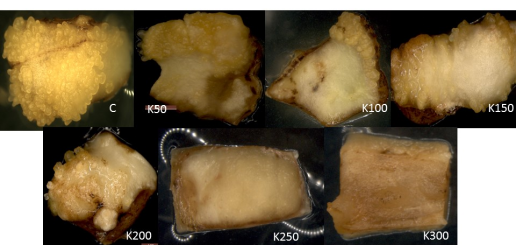

**Figure S1a.** Somatic embryos showing response over various concentrations of Kanamycin. C: Control, K50- K300: Kanamycin 50–300 mg/L.

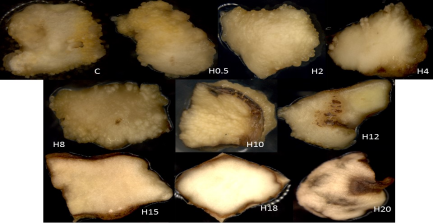

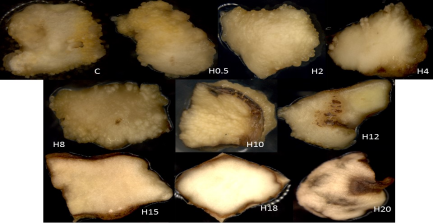

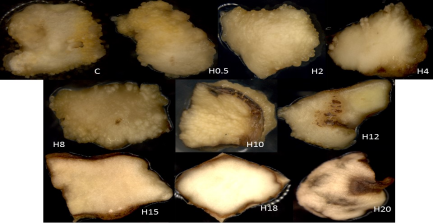

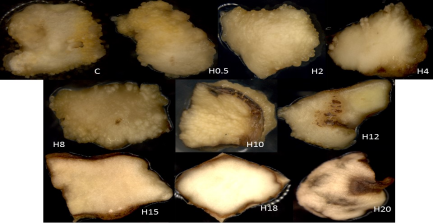

**Figure S1b.** Somatic embryos showing response over various concentrations of Hygromycin; c: Control, H0.5 - H20: Hygromycin 0.5-20 mg/L.

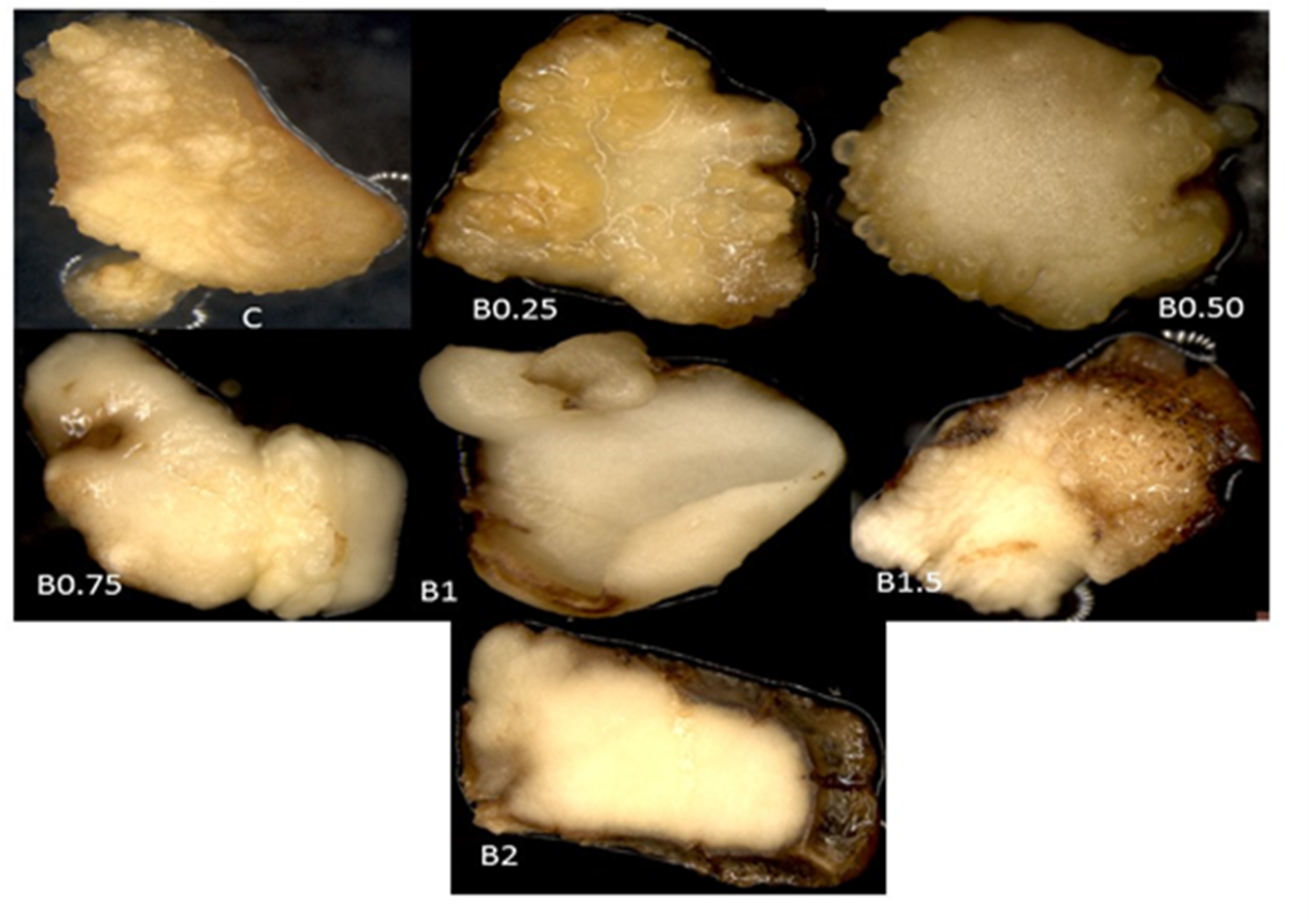

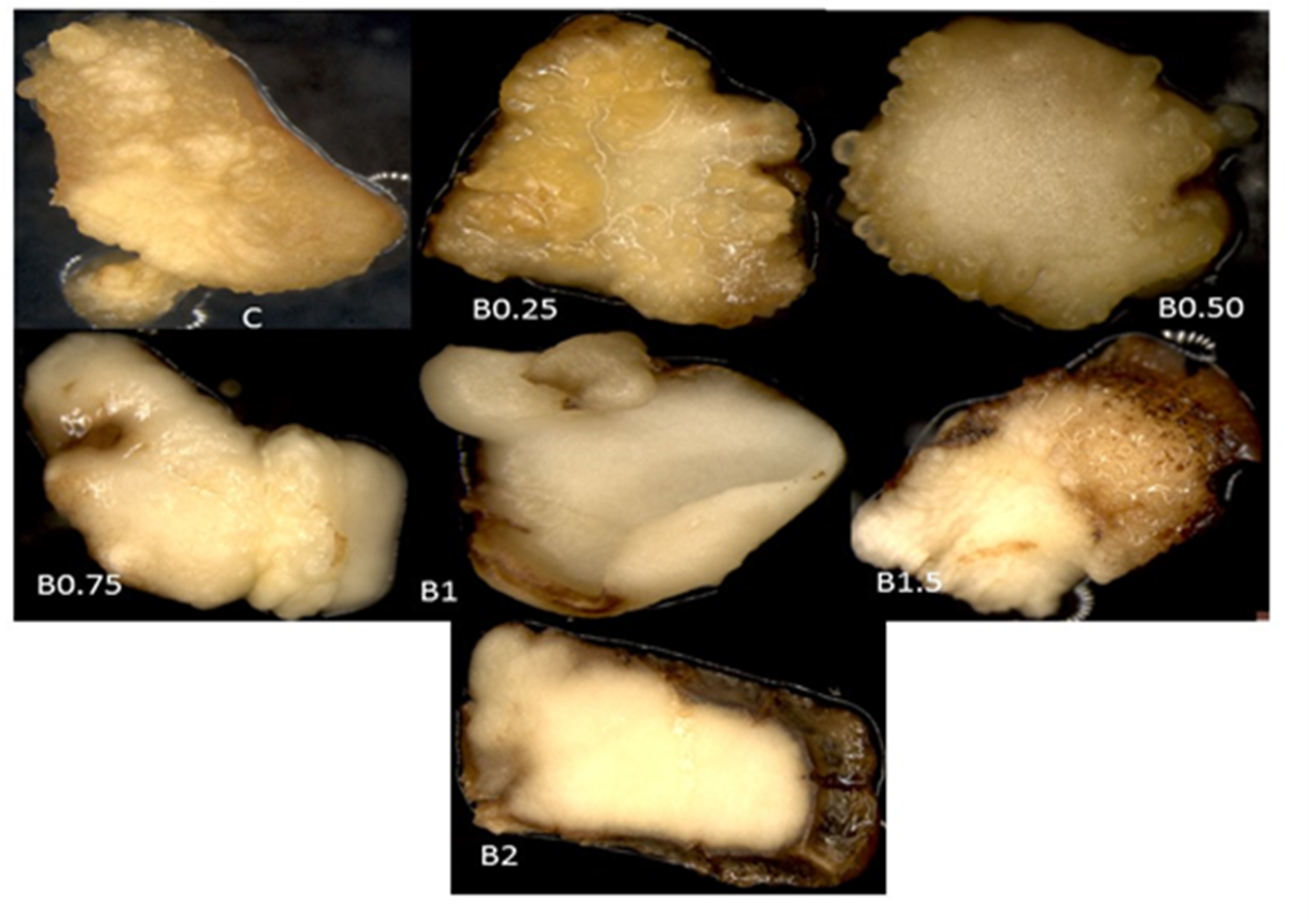

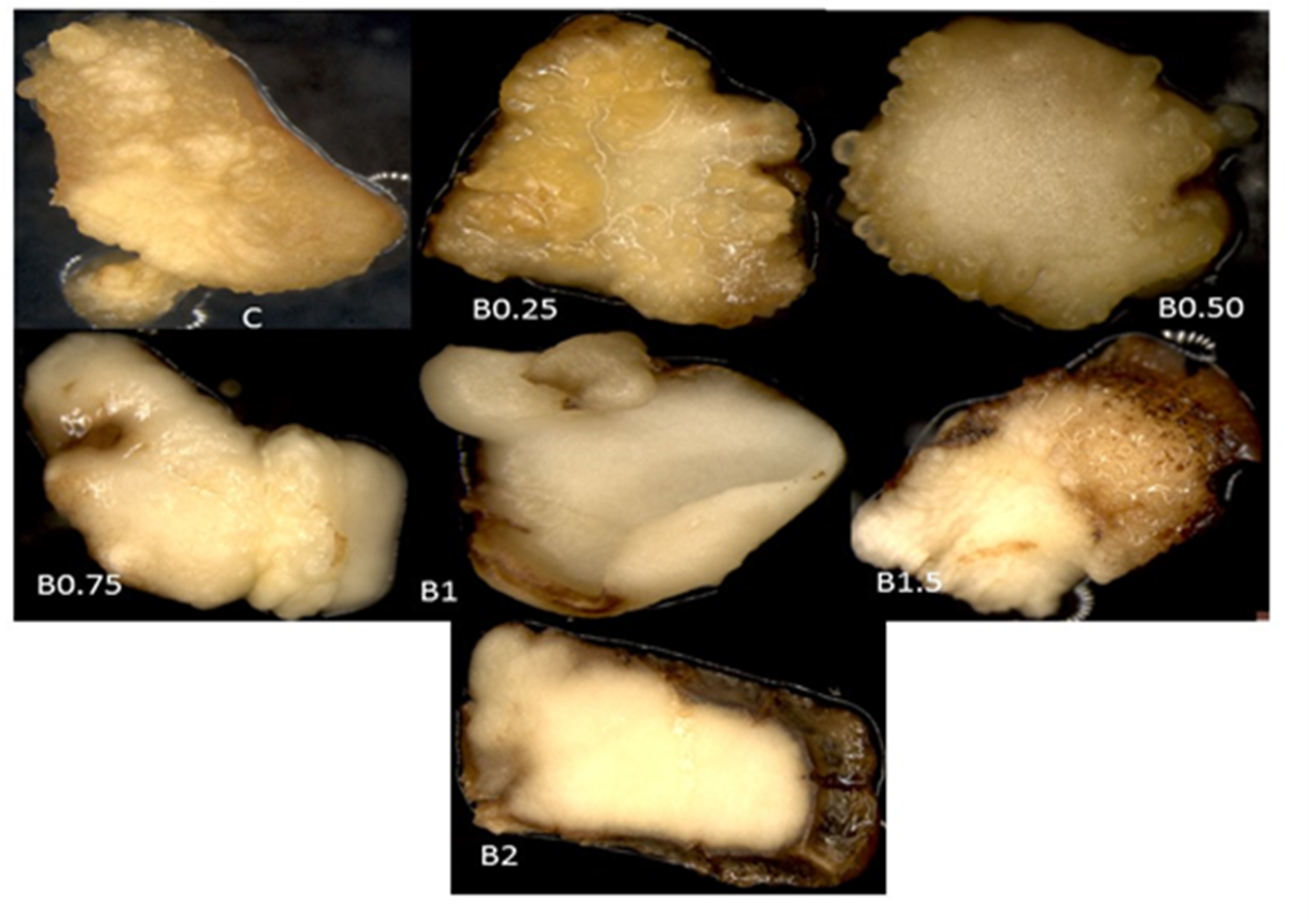

**Figure S1c.** Somatic embryos showing response over various concentrations of BASTA; C: Control, B0.25 – B2: BASTA 0.25 - BASTA2mg/L
