## Supplementary figures and images for "An optimized somatic embryo transformation system assisted homozygous edited rubber tree generation method mediated by CRISPR/Cas9"

### Supplemental Figure 2

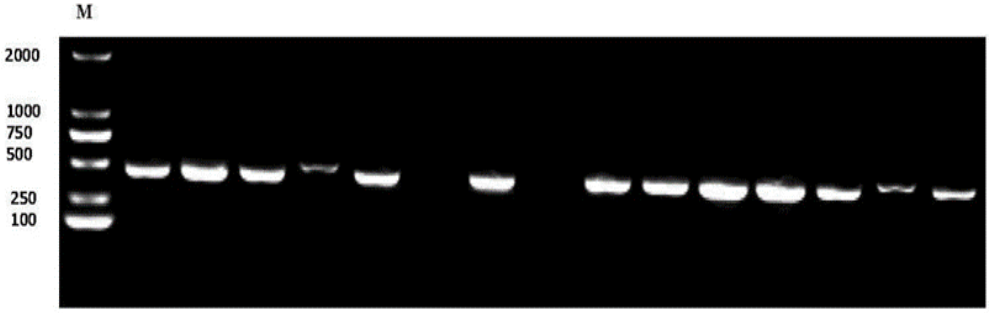

### Supplemental Figure 3

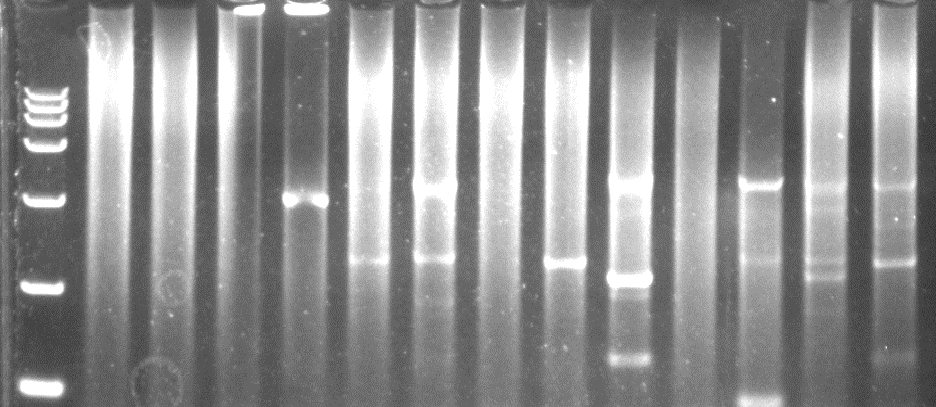

### Supplemental Figure 4

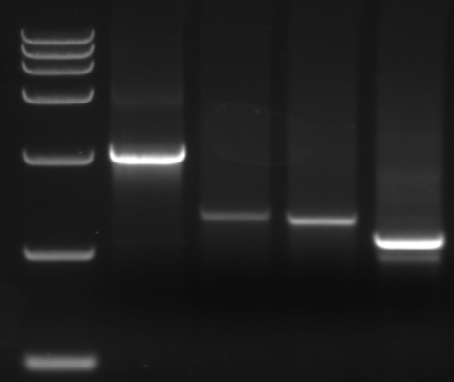
