## Supplemental Figure 5 for "An optimized somatic embryo transformation system assisted homozygous edited rubber tree generation method mediated by CRISPR/Cas9"

**①**

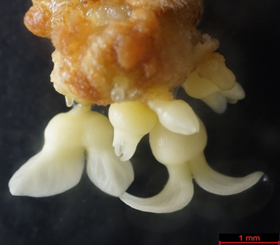

**④**

**③**

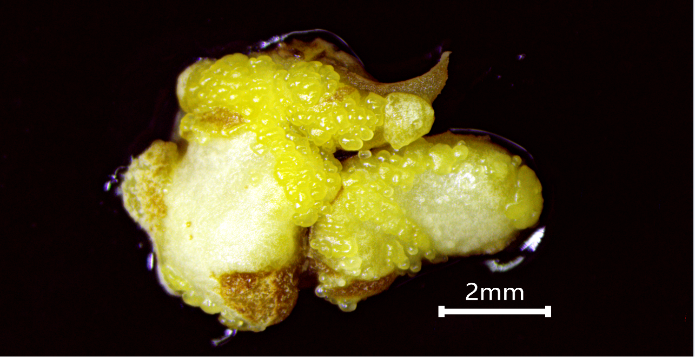

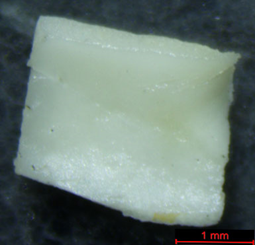

**②**

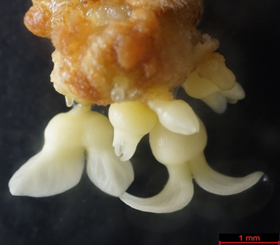

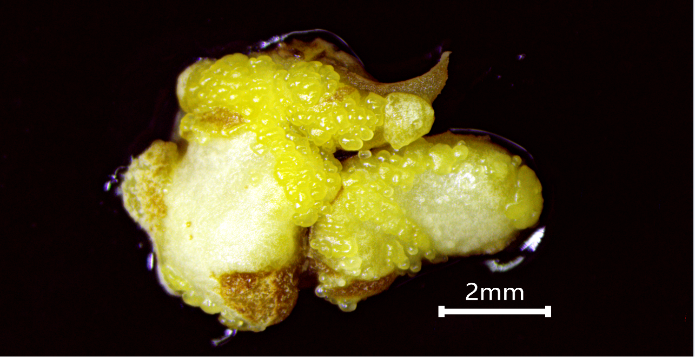

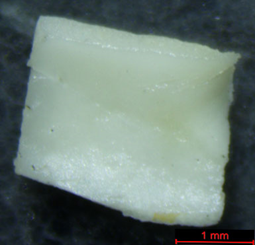

**⑧**

**T1（Homozygous**

**edited embryos）**

**⑦**

**⑤**

**⑥**

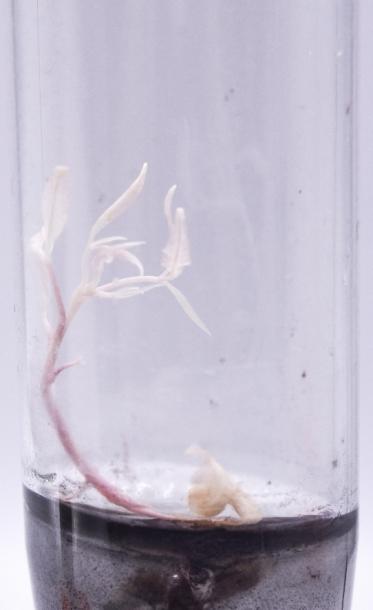

**Embryogenesis**

**Plant regeneration**

**Edited embryos are sliced**

**T0（Chimeric and**

**low editing ratio）**

**Embryogenesis**

**Callus**

**induction**

**Cutting**

**embryo**

**Callus**

**induction**

**Molecular detection**

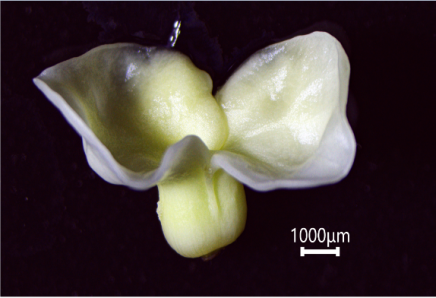

***Agrobacterium* infection**
